# Distribution and environmental drivers of multi-functional community traits in the Sundarbans mangrove forest

**DOI:** 10.1101/2025.07.15.664870

**Authors:** Tanjena Khatun, Md Rezaul Karim, Mohammed A.S. Arfin-Khan, Nabanita Karmaker, Md. Shamim Reza Saimun, Md Shydul Amin, Sanjeev K. Srivastava, Sharif A. Mukul, Fahmida Sultana

**Affiliations:** Department of Biological Science, University of Quebec in Montreal (UQAM), 141 Avenue President-Kennedy, Montreal, QC H2X 1Y4, Canada; Institute of Forestry and Conservation, John H. Daniels Faculty of Architecture, Landscape, and Design, University of Toronto, 33 Willcocks Street, Toronto, ON M5S 3B3, Canada; Department of Forestry and Environmental Science, School of Agriculture and Mineral Sciences, Shahjalal University of Science and Technology, Sylhet 3114, Bangladesh; School of Science Technology and Engineering (SSTE), University of the Sunshine Coast, Maroochydore DC, Queensland 4556, Australia; Department of Environment and Development Studies, United International University, Dhaka 1212, Bangladesh; Department of Earth and Environment, Florida International University, Miami, FL 33199, USA; School of Biosciences, Geography and Physics, Faculty of Science and Engineering, Swansea University, Singleton Park Swansea SA2 8PP Wales, UK

**Keywords:** Mangrove, Salinity gradient, Functional traits, Sundarbans, Functional group

## Abstract

Mangrove forests are vital blue-carbon ecosystems, playing a crucial role in global carbon storage, climate regulation, and biodiversity conservation. However, escalating environmental threats, particularly salinity intrusion, necessitate a deeper understanding of their functional dynamics to inform effective conservation and restoration strategies. This study examines the distribution of multi-functional community traits (MFCT) and their environmental drivers across mangrove functional groups (pioneer, mid-successional, climax) along salinity gradients in the Sundarbans, the world’s largest mangrove forest. Using data from 62 plots, nine functional traits from 17 mangrove species were analysed to evaluate MFCT diversity and its environmental determinants. Results revealed distinct variations in MFCT diversity, with mid-successional species exhibiting the highest trait diversity (64.0 ± 1.58), followed by climax (51.3 ± 2.09) and pioneer groups (50.2 ± 2.26). Climax species demonstrated resilience to salinity increases (R² = 0.29, p < 0.001), whereas mid-successional and pioneer groups were negatively affected (R² = 0.22, p < 0.001 and R² = 0.17, p < 0.001). Annual precipitation exerted contrasting effects, reducing diversity in climax species (R² = 0.25, p < 0.001) but enhancing it in mid-successional (R² = 0.22, p < 0.001) and pioneer groups (R² = 0.06, p = 0.05). Notably, MFCT values remained stable across salinity gradients, underscoring the ecosystem’s functional resilience. These findings highlight mid-successional species as keystones in sustaining mangrove multifunctionality and resilience under environmental stressors. Prioritizing these species in conservation and restoration efforts can mitigate the impacts of salinity intrusion and climate variability. By integrating the MFCT framework, resource managers can develop targeted restoration strategies to enhance ecosystem functionality and long-term stability. These insights provide a critical foundation for policymakers to align global mangrove conservation with adaptive management strategies, reinforcing coastal resilience, biodiversity conservation, and climate change mitigation.

## INTRODUCTION

Mangrove forests are critical blue-carbon ecosystems, renowned for their exceptional capacity to store carbon and mitigate global warming [1]. Beyond carbon storage, these forests play a pivotal role in maintaining biodiversity across terrestrial and marine ecosystems [2]. Despite their ecological and socio-economic significance, mangrove biodiversity is declining at an alarming rate [3]. Globally, mangroves are among the most threatened ecosystems, with a 3.4% reduction in mangrove area reported from 1996 to 2020 due to natural and anthropogenic disturbances [4]. Among these threats, salinity intrusion driven by sea-level rise poses a significant challenge, exacerbated by global warming and shifts in temperature and precipitation patterns [5].

To cope with these challenges, mangroves exhibit specialized morphological, physiological, and ecological traits enabling survival in extreme environments, including high salinity, strong winds, elevated temperatures, siltation, and oxygen-deprived soils [6]. These eco-physiological adaptations confer resilience to environmental stressors [7,8]. Within mangrove ecosystems, species are distributed across functional groups, each exhibiting unique trait combinations that enable adaptation to specific environmental gradients [9]. However, our understanding of how these traits respond to varying environmental stressors and which functional groups provide the greatest resilience remains limited. Additionally, knowledge gaps persist regarding how functional trait composition across groups shapes overall ecosystem functioning [10].

Biodiversity is a key driver of ecosystem functioning in forest ecosystems, including mangroves (Rahman et al., 2021). The insurance hypothesis suggests that biodiversity buffers ecosystem functioning against environmental stress, ensuring stability and resilience under changing conditions [12]. Similarly, the stress gradient hypothesis posits that species interactions shift from competition under favourable conditions to facilitation in stressful environments [13]. Such facilitation enhances complementarity effects among species, leading to greater resource use efficiency and ecosystem stability [14]. In mangrove ecosystems, including the Sundarbans, stress- tolerant species often dominate under adverse conditions such as salinity intrusion, disproportionately influencing ecosystem processes and maintaining critical functions [15]. It is important to note that, while these hypotheses are generally applicable, mangrove forests may exhibit exceptions to broader ecological patterns, with functional diversity potentially varying between pioneer, mid-successional, and climax species depending on specific environmental conditions.

To deepen our understanding of these dynamics, trait-based approaches have emerged as a powerful framework for understanding the relationship between biodiversity and ecosystem functioning [16–18]. Functional traits—encompassing morphological, biochemical, physiological, structural, phenological, and behavioural characteristics—provide a mechanistic link between biodiversity and ecosystem processes [19]. However, predicting community and ecosystem responses to environmental changes remains challenging due to the complexity and context- specificity of trait-environment relationships [20]. To address this, species can be grouped into functional groups based on shared traits and ecological roles, simplifying complexity and enhancing predictive capacity [21].

The Sundarbans, recognized as the world’s largest contiguous mangrove forest and a UNESCO World Heritage Site, play a crucial role in supporting coastal economies and maintaining global environmental balance. [22]. Species in the Sundarbans are categorized into pioneer, mid- successional, and climax groups based on their tolerance to sediment dynamics and elevation gradients [23]. Rising salinity, a consequence of sea-level rise, poses a significant threat, with projections indicating a 50% increase by 2050, potentially reducing forest productivity by 30% [24]. While previous studies have examined trait-based responses in the Sundarbans using single-and multi-trait approaches [24,25,23], the role of functional groups in shaping community responses and their influence on ecosystem multifunctionality remains understudied. To bridge this gap, we propose an integrated multi-trait approach termed Multi-Functional Community Traits (MFCT), inspired by the concept of ecosystem multifunctionality [26]. This framework aims to comprehensively assess the distribution, variation, and environmental drivers of functional traits across mangrove functional groups. Our study tests three hypotheses: (1) Trait diversity increases from pioneer to climax stages during succession; (2) Significant variation exists in MFCT diversity across functional groups and salinity levels; and (3) Among environmental factors, salinity exerts the most significant impact on MFCT diversity across functional groups.

## METHODS

### Study area

The Sundarbans Mangrove Forest, situated along the northern coast of the Bay of Bengal, spans between 89°00′ to 89°55′E and 21°30′ to 22°30′N [27]. Approximately 60% of this globally significant ecosystem lies within Bangladesh, comprising 40% of the country’s forested area, while the remaining 40% extends into India [28]. Designated as a UNESCO World Heritage Site, the Bangladeshi Sundarbans is divided into four ranges and 55 compartments. Elevations generally range between 0.5 and 3.0 meters above sea level (asl), with nearly 70% of the area below 1 meter [29]. Salinity is the dominant environmental driver shaping species composition in the Sundarbans. Salinity levels exhibit seasonal and spatial variations, increasing from east to west during the monsoon and from northeast to southwest in other seasons, peaking between April and May [29,30]. Based on salinity gradients, the forest is divided into three zones: freshwater (0.5–5 ppt during the monsoon), moderate saline (5–18 ppt), and highly saline areas with soil salinity exceeding twice that of freshwater zones [31]. The eastern and northeastern regions benefit from freshwater inflows from the Ganges, while the middle section is moderately saline with elevations of 3–4 meters asl. The southern and southeastern zones are characterized by high salinity levels [32]. The forest has a rich mixture of flora, fauna, and complex ecosystem functions, which makes it a unique ecosystem in the world [25].

### Sampling design & data collection

A total of 62 sampling plots (20 m × 20 m) were established across 54 management compartments of the Sundarbans using a random sampling strategy. Field data were collected from October 2021 to March 2022. The geographical coordinates of each plot were recorded using a handheld GPS (GPSMAP-65, Garmin Ltd., Olathe, KS, USA). All tree species within each plot were identified following the "Handbook of the Selected Plant Species of the Sundarbans and the Embankment Ecosystem" [33]. In total, 17 tree species were identified across the plots Mature leaves were collected from each plot to measure nine functional traits: chlorophyll content, leaf carbon content, leaf mass per area, leaf nitrogen content, leaf shape index, leaf succulence, specific leaf area, stomata size (guard cell length), and stomata density. On average, 3–5 leaves were sampled per tree for trait analysis. Due to field constraints in the soft, inundated soils of the Sundarbans, leaves were primarily collected from the lower canopy, which is the most accessible part of the tree. This methodology focused on shade leaves, as they are more easily reachable under these field conditions. To evaluate functional trait variation, mangroves were categorized into three functional groups—Pioneer, Mid-successional, and Climax—based on classifications from peer reviwed literature (Detailed sources are on Table 1)

**Table 1:** Mangrove species among different functional groups in the Sundarbans.

| Species | Functional group | Reference |
| --- | --- | --- |
| *Aegiceras corniculatum* | Pioneer | [45] |
| *Avicennia marina* | Pioneer | [46] |
| *Avicennia officinalis* | Pioneer | [47] |
| *Excoecaria indica* | Pioneer | [48] |
| *Kandelia candel* | Pioneer | [49] |
| *Rhizophora apiculata* | Pioneer | [50] |
| *Sonneratia apetala* | Pioneer | [46] |
| *Bruguiera gymnorrhiza* | Mid-successional | [46] |
| *Bruguiera sexangula* | Mid-successional | [46] |
| *Cerbera manghas* | Mid-successional | [51] |
| *Cynometra ramiflora* | Mid-successional | [52] |
| *Xylocarpus granatum* | Mid-successional | [53] |
| *Aglaia cucullata* | Climax | [54] |
| *Ceriops decandra* | Climax | [25] |
| *Heritiera fomes* | Climax | [25] |
| *Xylocarpus moluccensis* | Climax | [55] |
| *Excoecaria agallocha* | Climax | [25] |

### Measurement of plant functional traits

We selected 9 functional traits to represent the major axes of plant functional variation in this study. We measured chlorophyll content (mg/g tissue), leaf carbon content (%), leaf mass per area (LMA) (g/cm^2^), leaf nitrogen content (mg/g), leaf shape index (LSI), leaf succulence (LS) (g/cm^2^), specific leaf area (SLA) (cm^2^/g), Stomata size (guard cell length) (µm) and density (number/mm²). Details of all measurement laboratory methods are provided in Supplementary material 1.

### Soil sampling and analyses

Soil samples were collected from two subplots within each plot to a depth of 100 cm, subdivided into five layers: 0-10 cm, 11-20 cm, 21-30 cm, 31-50 cm, and 51-100 cm. Samples from the subplots were pooled before analysis. Detailed methods for soil sample analysis are provided in Supplementary material 2.

### Environmental factors

Electrical conductivity (EC) was used as a direct indicator of salinity. Saline zones were classified according to EC values: low salinity was defined as EC below 3.50 mS/cm, medium salinity ranged from 3.50 to 5.50 mS/cm, and high salinity was identified by EC values above 5.50 mS/cm. Mean annual temperature (BIO1) and annual precipitation (BIO12) data have been collected from the World Climate Database at a resolution of 1 sq. km for our study area [34]. hese climate variables represent long-term averages derived from 30 years of data, specifically from the period 1970 to 2000. As no high-resolution climate datasets are available for more recent years, and given the high reliability of the WorldClim data, we used these long-term averages to assess climate conditions in our study area.

### Multi-functional community trait (MFCT) index

Multi-functional Community Trait (MFCT) index was calculated by first normalizing the trait values for each plot using the community-weighted mean (CWM) trait value. CWM represents the mean trait value of a community, weighted by the relative abundance of each species. It reflects the trait values of the most dominant species within the community and is calculated using the following equation (Eq. 1) [19]:

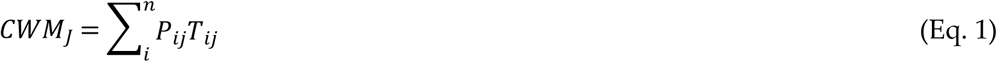

where P_ij_ is the abundance of the species i in the community j, and T_ij_ is the mean trait value of the species i in the community j.

After normalizing the trait values for each plot, a simple average was computed across the traits within each plot. This normalization process ensures that each plant trait contributes equally to the overall index, regardless of its natural variation or abundance within the community. By doing so, the MFCT index provides a robust, standardized measure (Eq. 2) that allows for meaningful comparisons of multiple traits across different plots and zones.

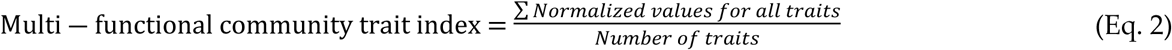

Higher values of the multi-functional community trait index indicate sites where functional trait diversity is higher across multiple dimensions.

### Statistical analysis

MFCT data from 62 plots were considered for spatial kriging in our study. Spatial dependence was assessed through variogram modelling, with parameters (sill, range, nugget, and fitted model) adjusted accordingly. Ordinary kriging was employed to interpolate these values across the study areas. Visualization of the results was performed using *‘ggplot2’*, with maps saved as PNG files. The entire workflow, encompassing variogram modelling, kriging, and visualization, was automated using a custom function based on the methodology outlined by [35].

All statistical analyses were performed using R version 4.4.1 (R Core Team, 2024). A linear model was chosen to explore the relationship between salinity and MFCT based on the assumption of linearity, providing a foundational understanding for subsequent analyses. Bar graphs with post hoc Tukey tests were generated using the *’ggplot2’* package, with Tukey’s Honestly Significant Difference (HSD) test applied through ’TukeyHSD’. Scatterplots and regression analyses were conducted using the linear model function (’*lm*’) in R. In addition, a Sankey diagram was constructed to visualize the relationships between salinity levels, plant traits, and functional groups, with data from 186 observations. This flow diagram employed a custom R script utilizing the *‘networkD3’* and *‘viridis’* packages, providing a clear representation of how traits and functional groups are influenced by salinity levels [36].

## RESULTS

### Spatial distribution of multi-functional community traits across functional groups

The spatial distribution of multi-functional community traits across pioneer, mid-successional, and climax species within the Sundarbans Mangrove Forest reveals distinct patterns along successional and environmental gradients (Figure 2). Climax species exhibit the highest multi-functional trait diversity, surpassing both mid-successional and pioneer species. This pattern indicates lower trait diversity in the early stages of mangrove succession, followed by a progressive increase as succession advances, culminating in peak diversity within climax communities. Pioneer species demonstrate elevated trait diversity in the northeastern region, where reduced salinity levels, driven by limited saltwater intrusion and increased freshwater influence, create favourable conditions. A secondary, albeit less pronounced, increase in trait diversity is observed in the southwestern region. Mid-successional species display a comparable spatial gradient but exhibit lower trait diversity in the southwestern areas.

**Figure 1:**
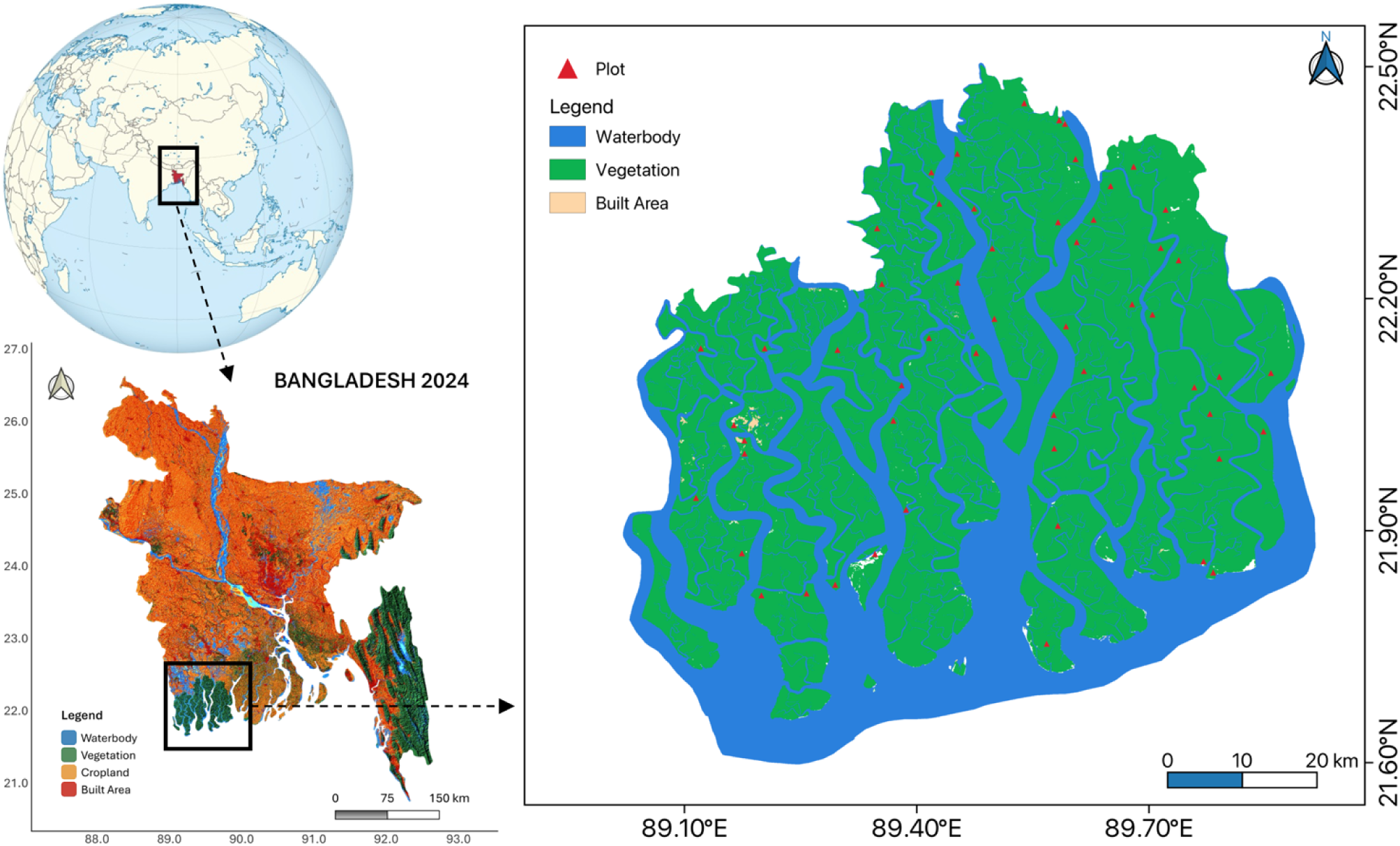
Map of the Sundarbans mangrove forest showing the location of study plots. The land use map of Bangladesh is based on the Sentinel-2 land use/land cover data for 2024.

**Figure 2:**
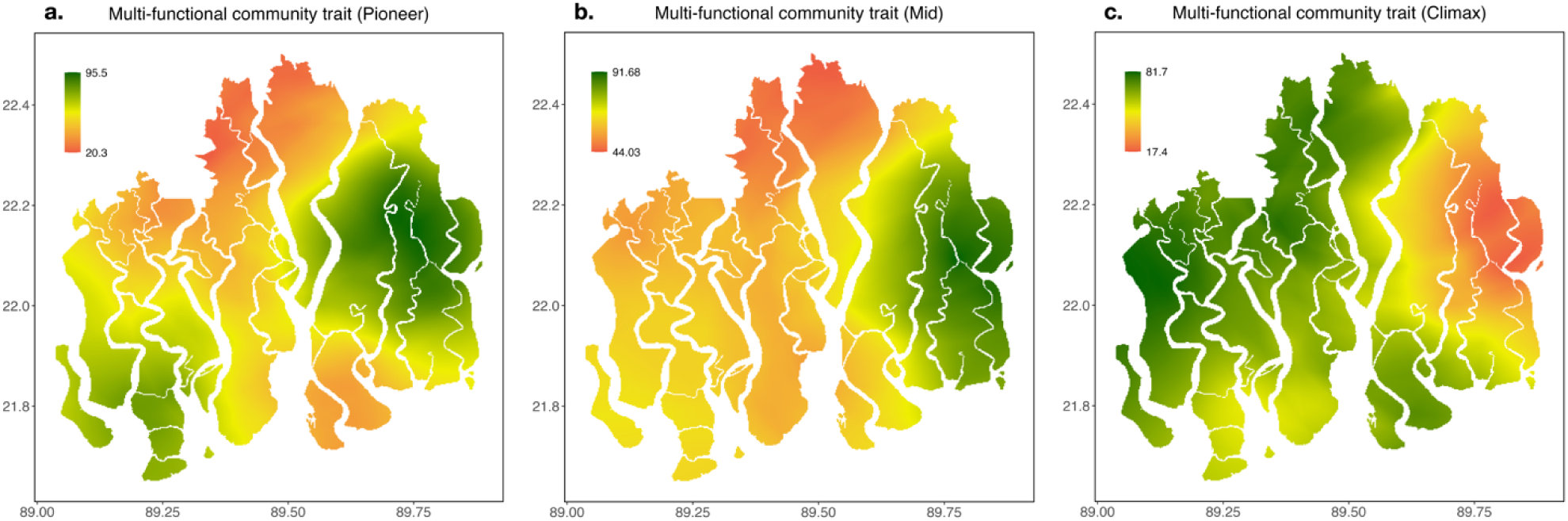
Kriging map depicting the distribution of multi-functional community traits across functional groups: (a) Pioneer, (b) Mid, and (c) Climax. The colour gradient from red to green represents increasing diversity in multi-functional community traits, with red indicating lower diversity and green indicating higher diversity

In contrast, climax species show consistently high trait diversity distributed across northern, southern, and southwestern regions. These areas, characterized by frequent tidal inundation and elevated salinity levels [37], appear to support a more functionally diverse assemblage of climax species, underscoring their resilience and adaptive capacity to environmental variability.

### Factors affecting MFCT trait across functional groups

Salinity and mean annual precipitation are significant drivers (p < 0.05) of MFCT across pioneer, mid-successional, and climax mangrove species (Figure 3). Climax mangroves exhibit a significant positive correlation with salinity (R = 0.77, p < 0.001), suggesting that elevated salinity enhances MFCT diversity in these species. In contrast, mid-successional and pioneer mangroves display significant negative correlations with salinity (R = -0.33, p < 0.001 and R = -0.34, p < 0.001, respectively), indicating that higher salinity suppresses trait diversity in these functional groups. While similar trends are observed with mean annual temperature, the correlations are not statistically significant (p > 0.05).

**Figure 3:**
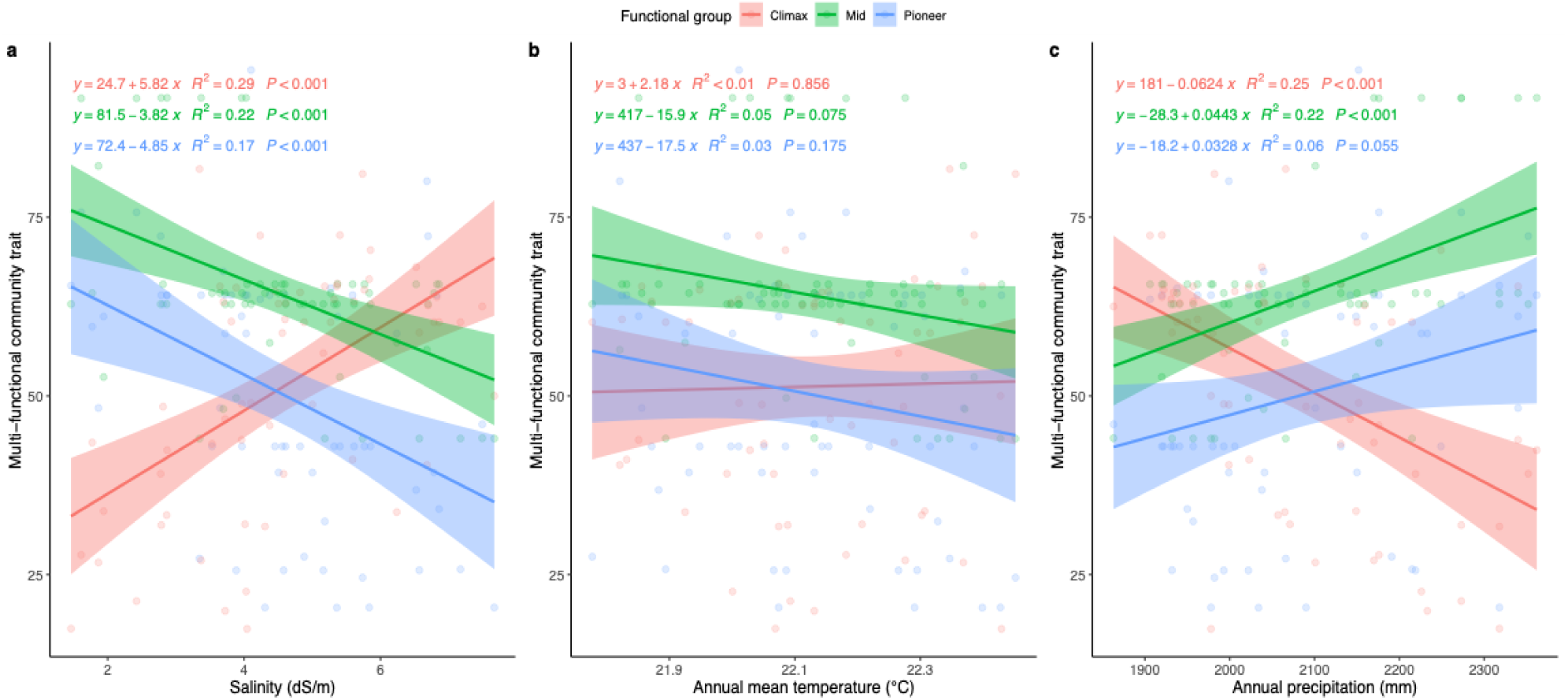
Line graphs showing the relationship between the multi-functional community trait and environmental variables for three functional groups of Pioneer, Mid, and Climax. The graphs illustrate how the multi-functional community trait varies with (a) salinity (dS/m), (b) annual mean temperature (°C), and (c) annual precipitation (mm).

Annual precipitation, however, demonstrates a pronounced influence on all three functional groups (p < 0.05). Climax mangroves show a significant negative correlation with precipitation (r = -0.49, p < 0.001), suggesting that increased precipitation reduces MFCT diversity. Conversely, mid-successional mangroves exhibit a positive correlation with precipitation (R = 0.29, p < 0.001), indicating an enhancement of trait diversity with higher precipitation levels. Pioneer mangroves also display a positive correlation with precipitation, though the association is comparatively weaker (R = 0.29, p < 0.05), implying that increased precipitation may have a modest effect on enhancing MFCT diversity in this group.

When considering the CWM traits individually (Figure 4), CWM Guard Cell Length is positively correlated with CWM Stomata Density (r = 0.65), which supports the linkage between stomatal traits in adaptation to salinity. CWM Leaf Nitrogen shows a negative correlation with CWM Leaf Carbon (R = -0.73), indicating a trade-off between these two traits in mangrove species. Notably, MFCT exhibits a strong positive correlation with CWM Stomata Density (R = 0.83), suggesting that species with higher MFCT diversity also tend to have larger or more abundant stomata, a trait linked to physiological responses to environmental stress.

**Figure 4:**
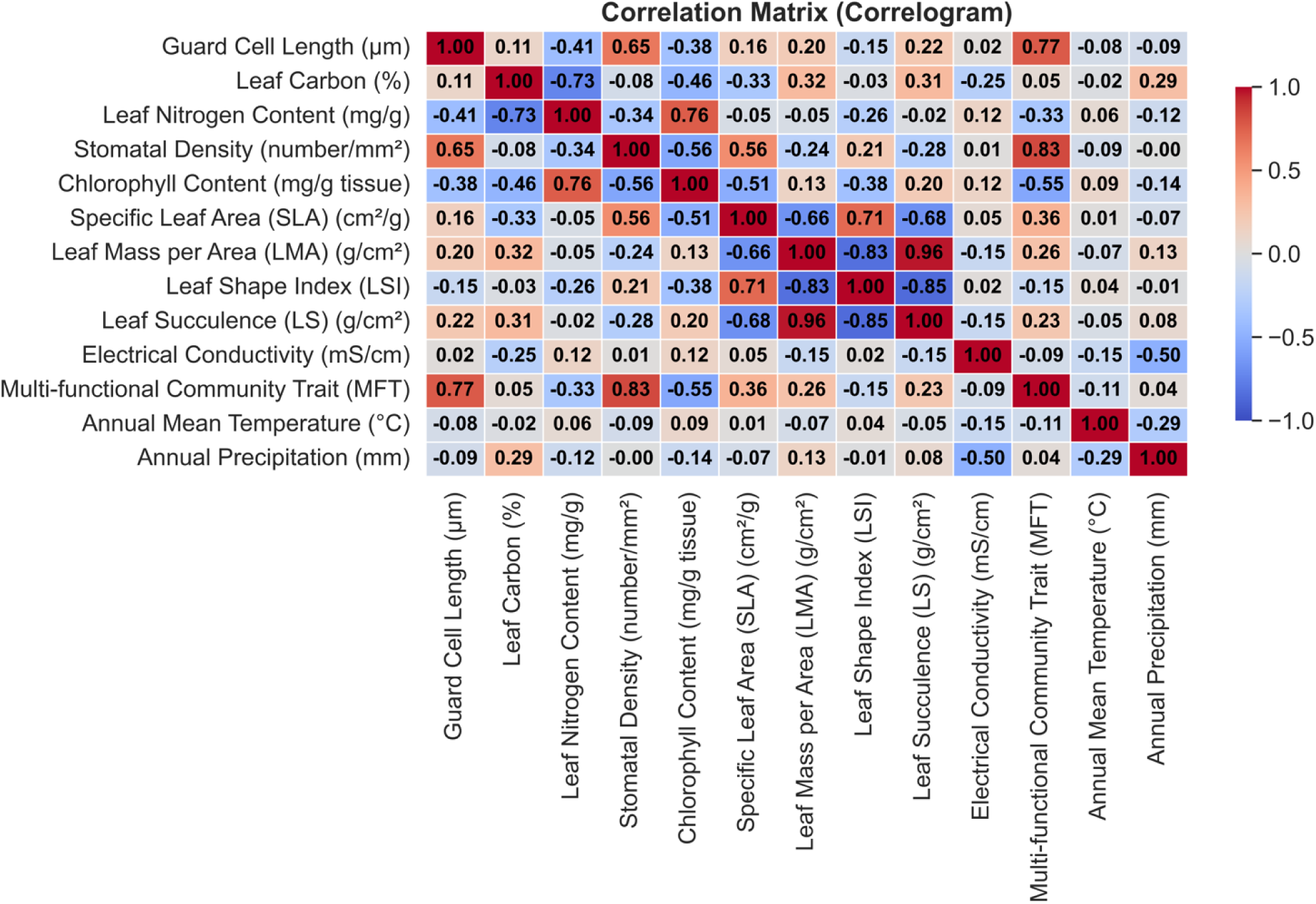
Correlation matrix of functional traits and environmental factors in the Sundarbans mangrove forest. The matrix shows relationships between leaf traits (e.g., guard cell length, leaf carbon, stomatal density), electrical conductivity (EC), multi-functional community trait (MFT), annual temperature (AMT), and precipitation (AP_mm). Correlation strength is represented by color intensity, with significant correlations highlighted in bold.

These findings underscore the divergent responses of functional groups to key environmental drivers, highlighting the contrasting ecological strategies adopted by pioneer, mid- successional, and climax mangroves under varying salinity and precipitation regimes.

### Comparison of MFCT across the functional groups and salinity levels

Multi-functional community trait (MFCT) values demonstrate significant variation (p < 0.05) across functional groups (Figure 5a). Among the three groups, mid-successional mangroves exhibit the highest MFCT values (64.0 ± 1.58, p < 0.05, marked with a), reflecting their greater trait diversity. In contrast, climax and pioneer mangroves exhibit lower MFCT values (51.3 ± 2.09 and 50.2 ± 2.26, respectively), with no statistically significant difference detected between these two groups (p > 0.05, marked with b). Across salinity gradients, MFCT values remain stable, with low (57.3 ± 3.48), medium (55.2 ± 1.58), and high (53.5 ± 2.14) salinity levels showing no significant differences (p > 0.05, marked with a; Figure 5b). These findings indicate that salinity exerts minimal influence on MFCT values, emphasizing the functional resilience of mangrove ecosystems to varying salinity levels.

**Figure 5:**
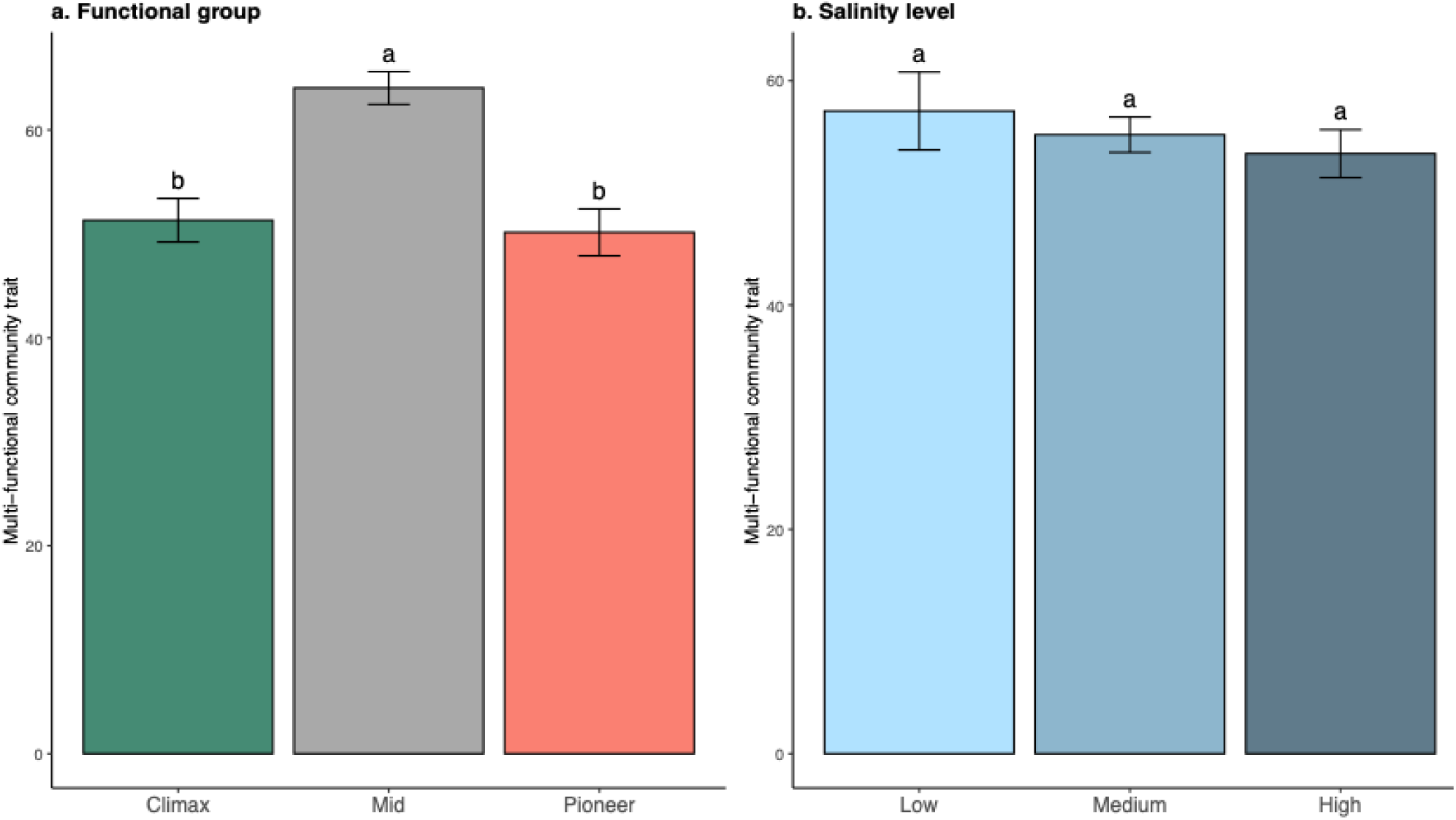
Bar plots depicting the distribution of the multi-functional community trait across different (a) functional groups and (b) salinity levels. Panel (a) compares the trait diversity among Pioneer, Mid, and Climax functional groups, highlighting differences in multi-functionality. Panel (b) illustrates the variation in the community trait across different salinity levels of low, medium, and high

The Sankey diagram provides further insight into the contributions of individual traits to these patterns (Figure 6). Notably, specific leaf area (flow values: high salinity: 173, medium salinity: 182, low salinity: 167) and stomata density (high salinity: 193, medium salinity: 189, low salinity: 190) emerge as dominant contributors to MFCT values across salinity gradients, underscoring their critical role in supporting ecosystem functionality and resilience. In contrast, traits such as total chlorophyll (high salinity: 0.000921, medium salinity: 0.000870, low salinity: 0.000811) and guard cell length (high salinity: 19.4, medium salinity: 19.1, low salinity: 18.6) display lower flow values, suggesting a relatively minor role in shaping overall MFCT values.

**Figure 6:**
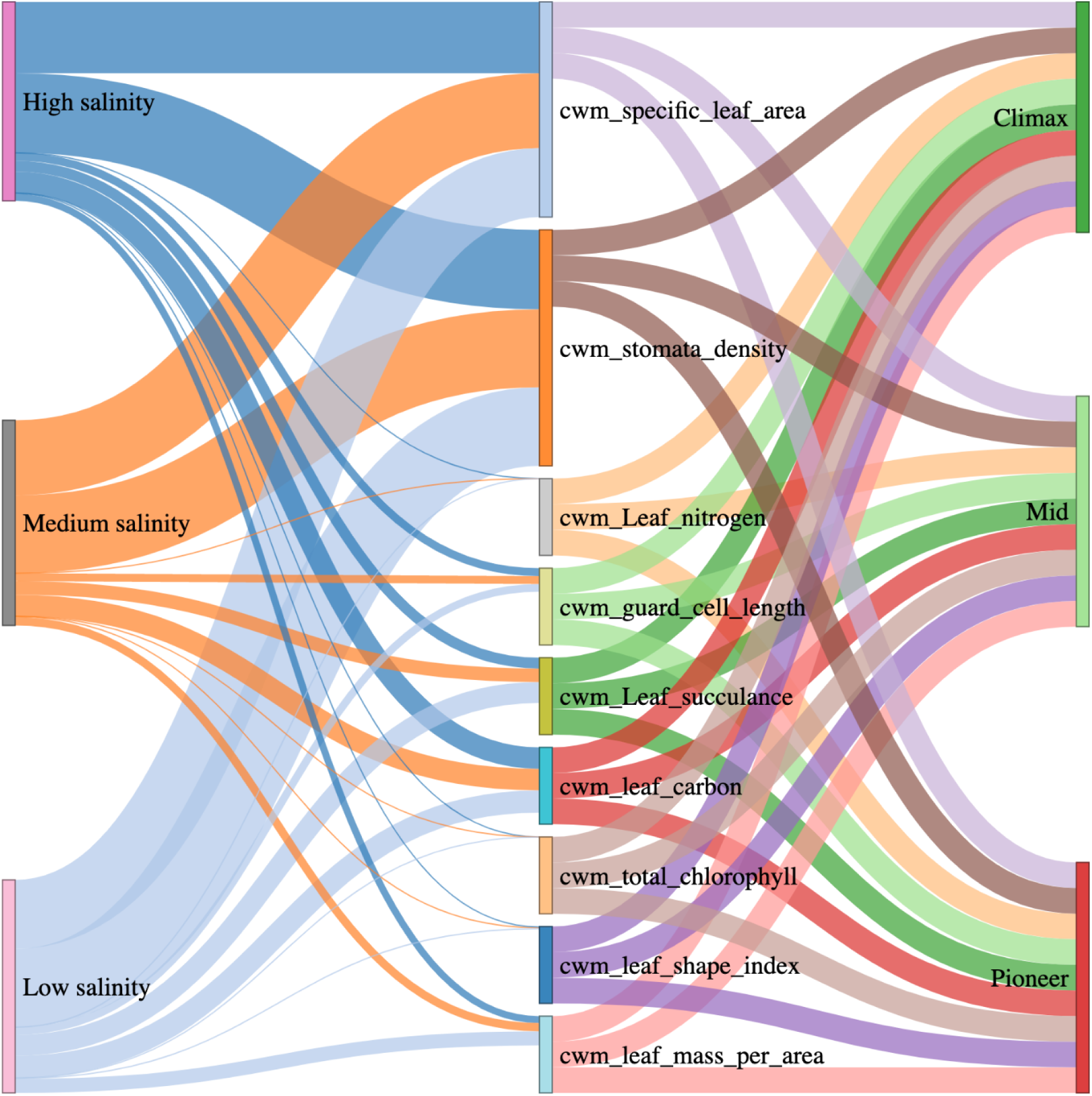
Sankey diagram illustrating the flow of individual trait contributions to multi-functional community trait (MFCT) values across salinity gradients and mangrove functional groups. Salinity levels (high, medium, and low) are shown on the left, community-weighted mean (CWM) functional traits in the middle of specific leaf area (SLA) (cm^2^/g), Stomata density (number/mm²), leaf nitrogen content (mg/g), Stomata size (guard cell length) (µm), leaf succulence (LS) (g/cm^2^), leaf carbon content (%), chlorophyll content (mg/g tissue), leaf shape index (LSI), leaf mass per area (LMA) (g/cm^2^) and functional groups (Pioneer, Mid, and Climax) on the right. The width of the flows represents the relative contribution of each trait

Traits like leaf succulence (high salinity: 25.9, medium salinity: 33.0, low salinity: 50.6) and leaf mass per area (high salinity: 16.8, medium salinity: 20.0, low salinity: 34.0) show moderate variability, suggesting localized contributions without overwhelming dominance. The analysis highlights the substantial influence of adaptive traits like specific leaf area and stomata density in driving ecosystem stability and resilience, even as other traits play more context-dependent roles. In summary, while trait diversity varies significantly across functional groups, the overall stability of MFCT values across salinity gradients reflects the robustness of mangrove ecosystems. The integration of statistical analysis and Sankey visualization underscores the centrality of key adaptive traits in maintaining ecosystem functionality and resilience across diverse environmental conditions.

## DISCUSSION

Our integrated multi-trait analysis highlights distinct spatial and functional patterns of multi- functional community trait (MFCT) diversity across functional groups in the Sundarbans Mangrove Forest Ecosystem. The observed increase in trait diversity from pioneer to climax species underscores the role of ecological succession in enhancing ecosystem functionality. Contrary to previous research suggesting higher diversity in low-salinity zones [24,25], our findings reveal that MFCT diversity at the community level is predominantly concentrated in moderate to high salinity areas.

### MFCT diversity across functional groups in the Sundarbans

Pioneer species, which endure extreme environmental stress during early succession, exhibit lower trait diversity due to survival-centric adaptations [38]. Over time, these species modify environmental conditions, facilitating the establishment of mid- and late-successional species [39]. Our findings reveal that climax species possess the highest spatial distribution and MFCT diversity, while mid-successional species exhibit the highest combined trait values. This indicates that mid-successional mangroves maintain a broader functional repertoire, which likely enhances ecosystem resilience.

Trait variation among species reflects diverse responses to environmental conditions. Coexisting species often show convergence or divergence in traits, shaping ecosystem functions. While trait convergence may lead to functional redundancy [40], it also reflects adaptive strategies that balance competition and functionality. These dynamics suggest that mid-successional mangroves, positioned between early colonizers and climax species, have evolved a wider range of functional traits, bolstering their ecological roles.

The spatial distribution of MFCT diversity supports our hypothesis that climax species exhibit the greatest diversity in moderate to high salinity zones. These regions, characterized by frequent tidal inundation and elevated salinity, present challenging conditions that limit site quality [41]. Despite this, climax mangroves thrive, suggesting their capacity to adapt and maintain functional diversity under stress. Conversely, less saline zones with substantial freshwater input favor late-successional species due to improved site quality [24]. Historical observations linked the highest mangrove diversity to low-salinity areas [24]. However, shifts in species composition, particularly in high-salinity zones, reflect the growing dominance of climax species like *Ceriops decandra* and *Excoecaria agallocha* [24,25]. Our findings align with these patterns, demonstrating that increased trait diversity among climax mangroves is an adaptive strategy that enhances resilience to adverse environmental conditions.

Trait diversity in pioneer and mid-successional mangroves is more pronounced in low-salinity zones, particularly in the northeastern Sundarbans. This aligns with ecological expectations for mid-successional species, which are less tolerant of salinity and thrive in freshwater-dominated areas [29,33]. Interestingly, pioneer species also exhibit slightly higher diversity in the southwestern region, despite its high salinity. This suggests that factors beyond salinity, such as local ecological interactions or resource availability, may influence pioneer species’ diversity. These patterns underscore the adaptability of mangrove communities across successional stages. By exhibiting differential responses to environmental gradients, mangroves contribute to the stability and functionality of the ecosystem under varying conditions.

### Drivers of MFCT in Sundarbans mangroves

Salinity and annual precipitation emerge as key factors shaping MFCT diversity across pioneer, mid-successional, and climax mangroves, corroborating our second hypothesis. Salinity positively influences climax species, supporting greater MFCT diversity. This trend aligns with previous findings [25], which attribute increased diversity to the dominance of highly salt-tolerant climax species such as *Excoecaria agallocha* and *Ceriops decandra*. Conversely, salinity negatively impacts the diversity of pioneer and mid-successional species, indicating that higher salinity limits their functional traits.

Precipitation also plays a complex role, exerting contrasting effects across successional stages. While it negatively affects climax mangroves, likely due to increased inundation and nutrient alterations [42], precipitation enhances trait diversity in mid-successional and pioneer species. These differential responses suggest that precipitation serves as a crucial ecological driver, influencing the functional diversity of mangrove communities in distinct ways.

### Variation in MFCT across functional groups and salinity gradients

Significant variation in MFCT diversity across functional groups highlights the adaptive strategies of mangroves at different stages of ecosystem development. Mid-successional mangroves exhibit the highest MFCT values, reflecting their broad functional repertoire. Besides, the lack of significant difference between pioneer and climax mangroves suggests that despite occupying different niches in the successional process [43,44] both groups may have adapted to their specific roles in the ecosystem with comparable functional traits.

Interestingly, MFCT values remain relatively stable across salinity gradients. This finding, which contrasts with our third hypothesis, suggests that while salinity influences trait diversity within successional stages, the community as a whole maintains its functional capacity. Such stability indicates that mangrove species possess inherent resilience to salinity fluctuations, likely driven by their eco-physiological adaptations [7,8].

### Ecological implications and future research

Our results underscore the importance of trait-based approaches in understanding mangrove ecosystem dynamics. By revealing the drivers of MFCT diversity, this study highlights the adaptive strategies of mangroves across environmental gradients, providing insights into their resilience under changing climatic conditions. However, the mechanisms underpinning these adaptive responses remain poorly understood. Further research should focus on identifying the physiological and genetic factors that enable mangroves to thrive in diverse environments. Mangroves’ ability to maintain functional diversity despite environmental stressors positions them as critical components of coastal resilience strategies. Preserving their multifunctional traits is essential for sustaining ecosystem services, particularly in the face of rising sea levels and salinity intrusion. Our findings emphasize the need for conservation efforts that prioritize the protection of diverse mangrove communities, ensuring their continued role in supporting ecological and socio-economic systems.

## CONCLUSION

This study provides valuable insights into the spatial distribution, diversity, and drivers of multifunctional traits within the Sundarbans Mangrove Forest Ecosystem. Significant variation in trait diversity was observed across functional groups, with climax species showing the highest trait diversity in areas of moderate to high salinity, challenging previous species-specific studies that linked higher diversity to low salinity zones. Findings emphasize that salinity and annual precipitation are critical drivers shaping mangrove multifunctional traits, with salinity promoting diversity in climax species but negatively impacting pioneer and mid-successional species. The effect of precipitation is more complex, varying across successional stages. Despite variations in trait diversity, the overall mangrove community maintains resilience across salinity gradients, highlighting adaptive strategies to environmental fluctuations. However, the underlying mechanisms driving these responses remain unexplored, warranting further research to enhance the understanding of mangrove community dynamics. These findings are crucial for informing conservation strategies, habitat restoration, and prioritizing climate change resilience efforts globally, as mangroves continue to provide essential ecosystem services. The integrated multi-trait approach proposed here offers a robust framework for advancing mangrove ecology research and improving predictions of community responses to climate change.

## AUTHOR CONTRIBUTIONS

T. Khatun and M. R. Karim conceptualized the ideas and designed the methodology. M. R. Karim, M. A. S. Arfin-Khan, and S. A. Mukul collected the data, while T. Khatun and M. R. Karim led the data analysis. T. Khatun and M. R. Karim also co-led the writing of the original draft. T. Khatun contributed to software development and data curation, with M. R. Karim also contributing to software development. M. A. S. Arfin-Khan, N. Karmaker, M. S. R. Saimun, M. S. Amin, S. K. Srivastava, S. A. Mukul, and F. Sultana critically reviewed and edited the manuscript. Funding acquisition was led by M. A. S. Arfin-Khan, S. K. Srivastava, and S. A. Mukul. F. Sultana supervised the study. All authors critically contributed to the drafts and gave final approval for publication.

## ACKNOWLEGEMNETS

The authors would like to acknowledge the funding support received from the SUST Research Center (FES/2021/2/04, FES/2022/1/09, FES/2023/2/01, FES/2024/1/01) and Asia-Pacific Network for Global Change Research (APN, Japan; project code: CRRP2020-08MY-Srivastava). The British Ecological Society (UK; grant reference number: BES LRB20-1006), and the National Geographic Society (USA; grant reference number: NGS-78528R-22).

## DATA AVAILABILITY STATEMENT

All data that support the findings of this study are included in the article.

## SUPPLEMENTARY MATERIALS

### Supplementary material 1 Chlorophyll Content

Chlorophyll content was measured for each species using a spectrophotometer according to the procedure given by [56]. Three samples were prepared for each species using the freshly collected leaves. 80% Acetone was used as a solvent for the extraction of chlorophyll. Absorbance readings of chlorophyll extracts were measured at two different wavelengths 645 nm and 663 nm respectively. Based on the absorbance value, calculations were made using the equation suggested by Arnon’s (1949) and the amount of chlorophyll-a, chlorophyll-b and total chlorophyll were estimated.

The following formulas were used for calculating chlorophyll-a (Eq. 3), chlorophyll-b (Eq. 4), and total chlorophyll content (Eq. 5).

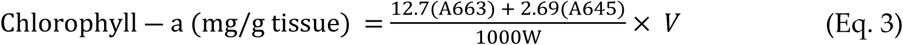

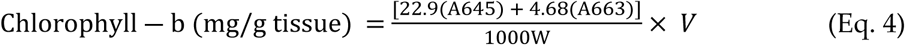

Here, A_645_ = Absorbance of TC extract at 645 nm, A_663_ = Absorbance of TC extract at 663 nm, V= Total extract volume, W= Leaf fresh weight

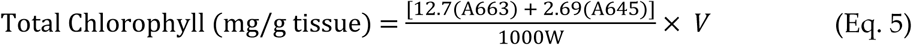

### Leaf Carbon Content

Estimation of leaf carbon content was done with the collected leaf samples. The collected leaf samples were first oven dried for 48 hours at 105°C to get the dried weight. Then the dried samples were grinded, and the grinded sample was taken in pre weighted crucibles. After that, the crucibles were placed in the maple furnace at 600°C temperature for two hours. The crucibles then cooled carefully inside the furnace. When crucibles with ash cooled down, the samples were weighted (Eq. 6), and percentage of leaf carbon calculated. Percentage of leaf carbon was calculated as Eq. 7

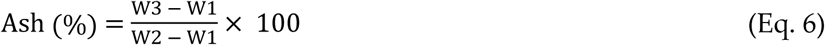

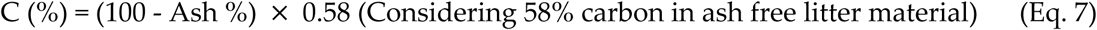

Here, C = Carbon, W1 = Weight of crucibles, W2 = Weight of leaf + Crucibles, W3 = Weight of ash + Crucibles

### Leaf Mass per Area

According to the protocols of [57], Leaf mass per area was measured. To measure leaf mass per area, leaf samples were put in the oven to dry for 72 hours in 70℃ temperature. The dry weight of the leaves was measured with a digital balance meter. Then the trait was calculated using the following (Eq. 8) formula:

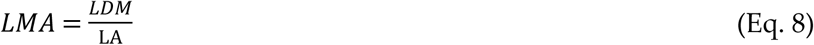

Here, LMA= Leaf Mass per Area (g/cm^2^); LDM = Leaf oven-dry mass (g); LA = Fresh leaf area (cm^2^)

### Leaf Nitrogen Content

Leaf nitrogen content was determined as nitrogen mass per gm tissue for the tree species found in the plot. The samples were first heated in the presence of conc. sulfuric acid, K_2_SO_4_ and CuSO_4_ in a Kjeldahl digestion apparatus and evaporated until SO3 fumes were obtained, and the solution became colourless or pale yellow. The residue was then cooled, diluted, treated, and made alkaline with a sodium-hydroxide solution. The ammonia in the sample is distilled and determined by titration after distillation. Total Kjeldahl nitrogen was then determined using the following formula (Eq. 9).

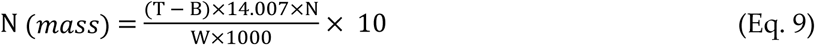

Here, T= amount of standard 2N H_2_SO_4_ solution used in titrating sample, B= amount of standard 2N H_2_SO_4_ solution used in titrating blank, N=normality of sulfuric acid solution, W= sample weight

### Leaf shape index

Leaf shape index is the ratio of leaf length and leaf width. For the calculation of leaf shape index, leaf length and width were measured using centimetre scale. The unit of leaf shape index is cm/cm [58].

### Leaf succulence

Leaf succulence was calculated according to [59] using the following equation (Eq. 10):

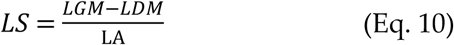

Here, LS= Leaf Succulence (g/cm^2^); LGM = Leaf green mass (g); LDM = Leaf oven-dry mass (g); LA = Fresh leaf area (cm^2^)

### Specific leaf area

Specific leaf area is the inverse of LMA (Leaf Mass per Area). This is expressed as cm^2^/g. The equation (Eq. 11) is as follows:

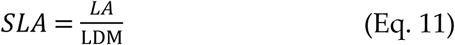

Where, SLA= Specific Leaf Area (cm^2^/g); LA = Fresh leaf area (cm^2^); LDM = Leaf oven-dry mass (g) [60]

### Stomata size and density

17 species identified in the plot were selected for measuring stomata size and density. However, stomata of some species were not identified despite using microscope, hence these species were excluded from the study and finally stomatal measurements were performed for 13 species representing 76% of the total individual found in the plot. Leaf samples were prepared with a transparent quick dry nail polish for stomatal measurements using nail polish imprint method [61,62] on the abaxial leaf surface [61,63], as in most species, stomata density is higher in the abaxial epidermis than in the adaxial epidermis.

Three microscopic slides were prepared as samples from each leaf by avoiding the major veins. Stomata were then observed by a digital light microscope. Two images were taken from each sample through the digital 8 MP scope mounted camera (MU800, Amscope, US). The number of stomata was counted in the image to calculate stomata density per mm². Guard cell length (GCL) was measured on the same image. A total of 30 stomata were randomly selected for a single image for the measurement of guard cell length.

Stomata density and guard cell length were then measured according to [64]. Thus, we measured abaxial stomata density on nine leaf samples with 18 images and measured GCL for 540 stomata for each species. The obtained measurements were then averaged to get stomata density and guard cell length of individual species.

### Supplementary material 2

Soil salinity was measured in the laboratory at the Department of Forestry and Environmental Science, Shahjalal University of Science and Technology, Bangladesh. Electrical conductivity (EC) was determined using a portable pH/EC/TDS meter (Model HI9810-61, Hanna Instruments, USA). A soil-to-distilled water ratio of 1:2 was used, with soil dilution applied to ensure accurate measurements, following the method outlined by [65]. The EC values for each of the five depth layers were averaged to provide a representative measure of soil salinity (mS/cm).

